## Supplemental Figures for "Alternative splicing controls teneurin-3 compact dimer formation for neuronal recognition"

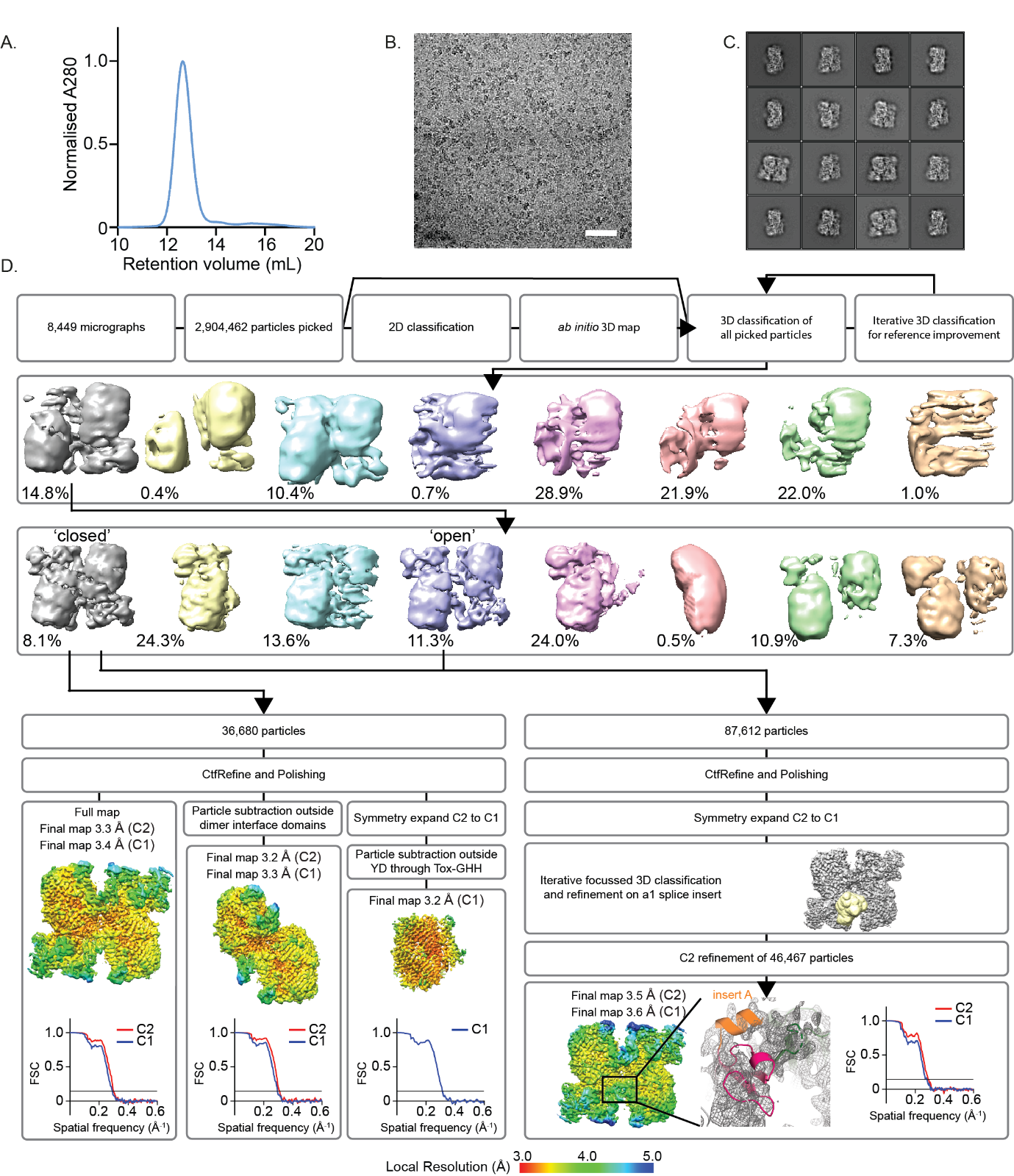

Supplemental Figure 1: Purification, cryo-EM data collection and reconstruction of teneurin-3 A_1_B_1_ compact dimer. **A)** Size exclusion chromatography (SEC) trace of the Ten3-A_1_B_1_ ectodomain after nickel affinity purification. **B)** Representative cryo-EM micrograph of vitrified Ten3-A_1_B_1_ ectodomain. Scale bar is 50 nm. **C)** Single-particle analysis (SPA) 2D classes of compact dimeric Ten3-A_1_B_1_ particles in ‘closed’ conformation. **D)** SPA workflow for the reconstruction of cryo-EM electron density of compact dimeric Ten3-A_1_B_1_. Left: reconstruction workflow of the full dimeric map. Second to left: dimeric map reconstruction after particles subtraction around interface domain. Third to left: C2-to-C1 symmetry-expanded reconstruction of subunit with density outside YD through Tox-GHH subtracted. Right: focussed classification and refinement of splice insert A (orange helix) with dimers from the closed conformation and more flexible open conformation. Corresponding local resolution values and Fourier shell correlation (FSC) graphs are shown for each map.

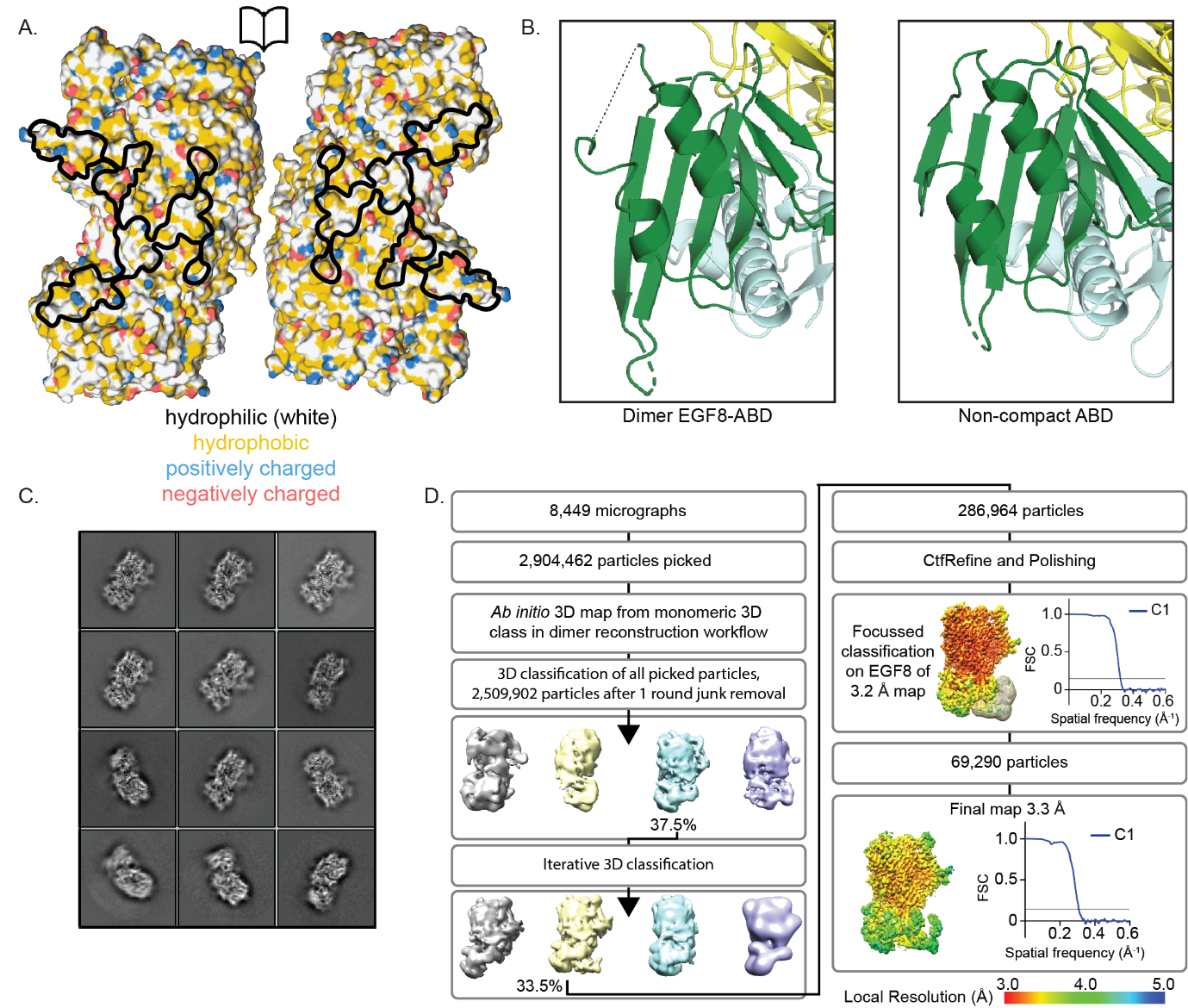

Supplemental Figure 2: Teneurin-3 A_1_B_1_ compact interface analysis and non-compact subunit reconstruction. **A)** Open book electrostatics and hydrophobicity surface representation of the Ten3 compact dimer. Black lines indicate the outline of compact dimer interface. **B)** Structural comparison of the Ten3-A_1_B_1_ non-compact subunit ABD versus the compact dimeric EGF8-contacting ABD. The non-compact ABD displays the presence of an additional β-strand at solvent-facing edge of the ABD β-sheet. Missing residues in the non-compact subunit are indicated with dashed black line. **C)** Single-particle analysis (SPA) 2D classes of Ten3-A_1_B_1_ non-compact subunit particles after focussed classification on EGF8. **D)** SPA workflow for the reconstruction of the Ten3-A_1_B_1_ non-compact subunit, including subsequent focussed classification and refinement of the EGF8-containing particles. Corresponding local resolution values and Fourier shell correlation (FSC) graphs are shown for each map.

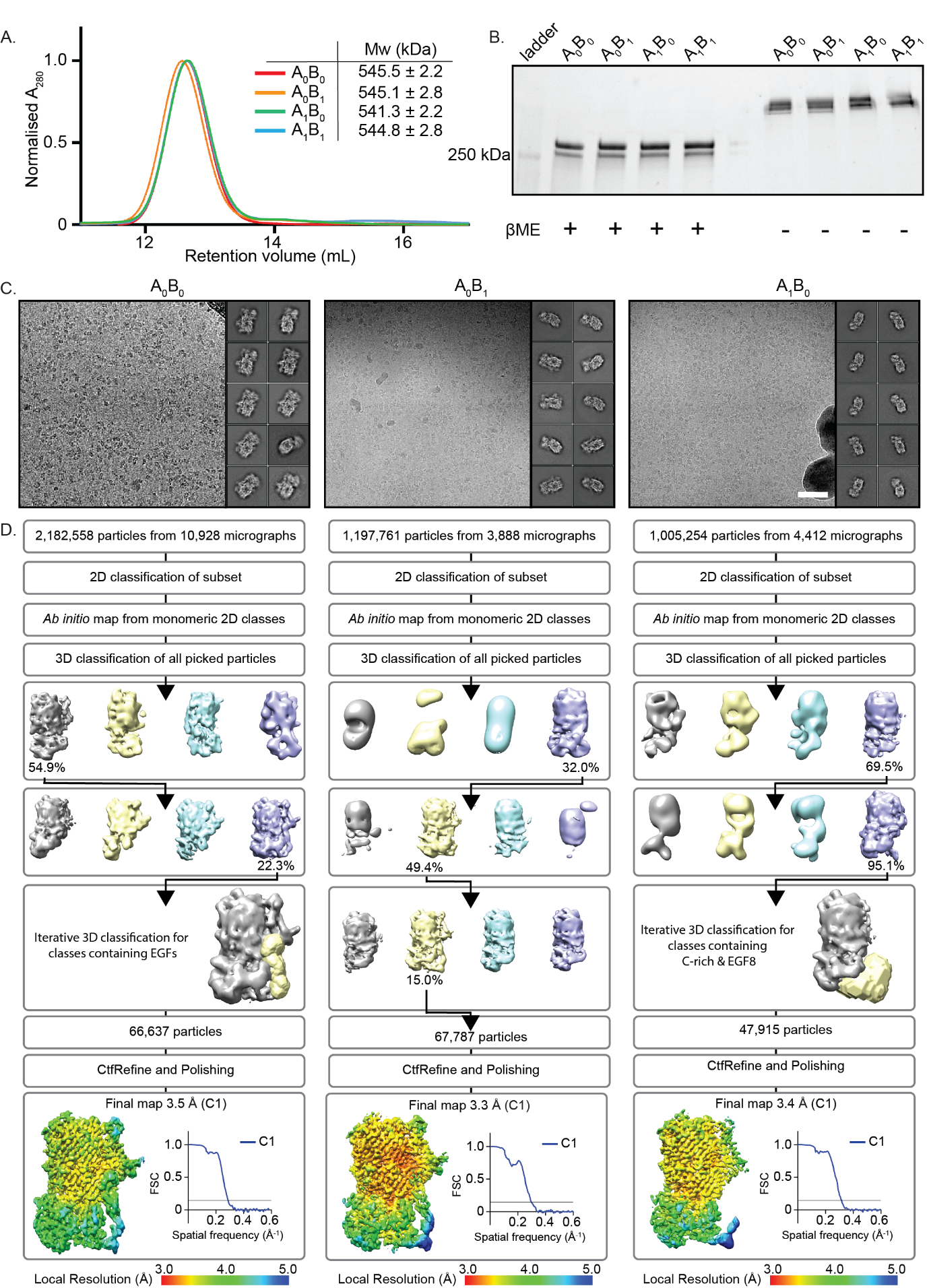

Supplemental Figure 3: Purification, cryo-EM data collection and reconstruction of teneurin-3 A_0_B_0_, A_0_B_1_, A_1_B_0_, non-compact subunits. **A)** Size exclusion chromatography (SEC) trace of all Ten3 ectodomains after nickel affinity purification. Inset displays multi-angle light scattering (MALS) data of all isoform directly after exiting the SEC column. **C)** PAGE gel of all purified Ten3 isoform ectodomain in the presence and absence of reducing agent β-mercaptoethanol (βME). **D)** Representative cryo-EM micrographs of vitrified Ten3 ectodomain and single-particle analysis (SPA) 2D classes for the non-compact A_0_B_0_, A_0_B_1_, and A_1_B_0_ isoform subunits. Scale bars are 50 nm. Per isoform 2D classes of the final reconstructions from single-particle (SPA) analysis in D are shown. **D)** SPA workflow for the reconstruction of cryo-EM electron density of Ten3 isoform subunits. Workflows include subsequent focussed classification and refinement of the EGF-containing particles. Corresponding local resolution values and Fourier shell correlation (FSC) graphs are shown for each map.

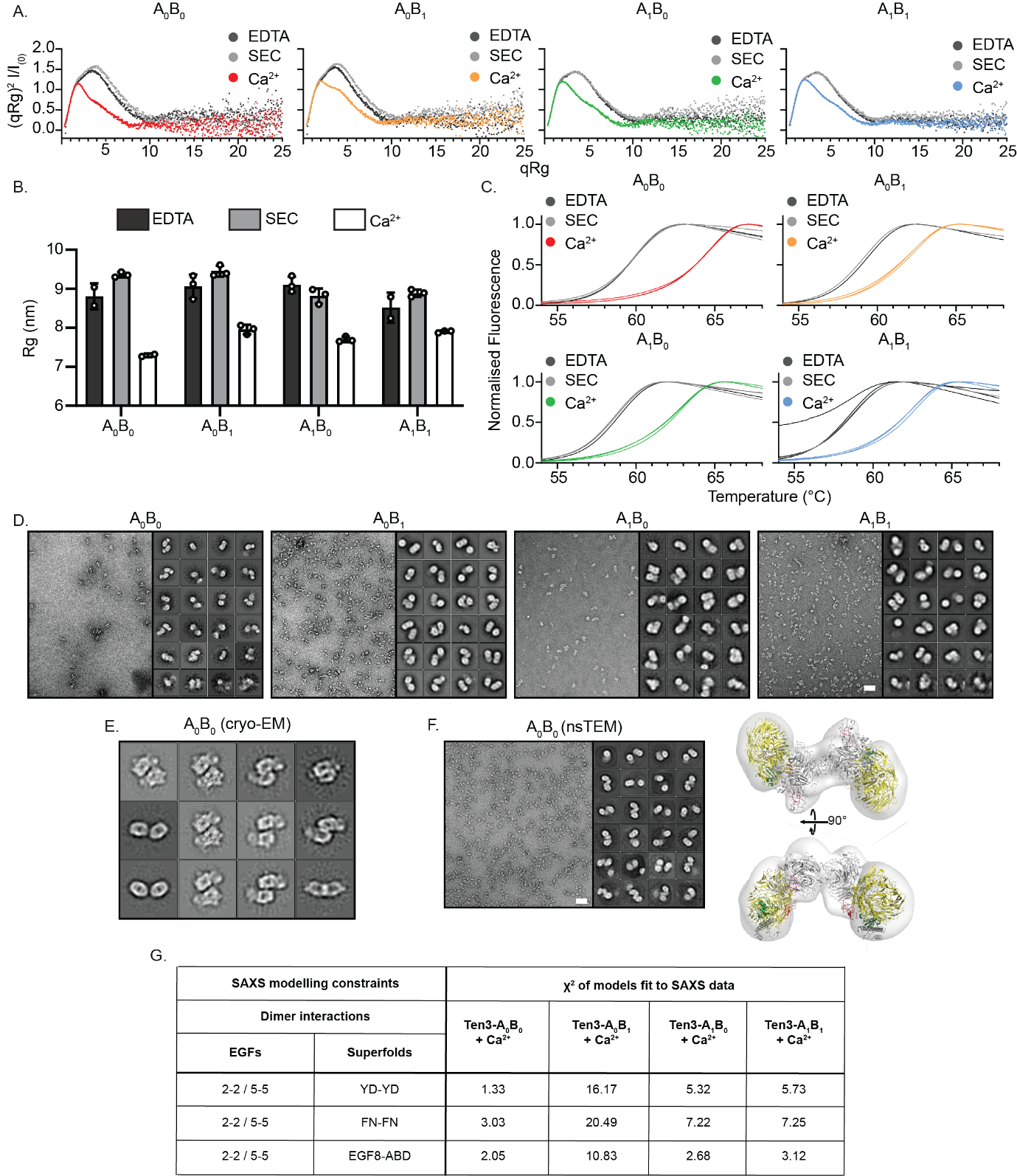

Supplemental Figure 4: Teneurin-3 isoform-specific compactness and stability assessment. **A)** Dimensionless Kratky-plots for the four purified Ten3 isoforms in the presence (Ca^2+^) and absence (SEC) of calcium, as well as under addition of EDTA (EDTA). **B)** SAXS-derived radii of gyration (Rg) in the presence and absence of calcium, as well as under addition of EDTA. Data points represent SAXS measurements repeats at Ten3 concentration of 1.0, 0.5, and 0.25 mg/mL. **C)** Thermal stability melting curves for the four purified Ten3 isoforms in the presence and absence of calcium, as well as under addition of EDTA. Per-condition triplicate experimental repeats are plotted. **D)** Representative negative stain EM micrographs of Ten3 ectodomain and single-particle analysis (SPA) 2D classes of all picked particles for each isoform indicated above. Scale bars is 50 nm. **E)** Cryo-EM SPA 2D classes of Ten3 A_0_B_0_ in a YD-YD-connected compact conformation. **F)** Negative stain TEM micrograph with SPA 2D classes and 3D refinement of the A0B0 NHL-NHL connected compact dimer. Scale bars is 50 nm. **G)** Fits of rigid-body models to SAXS data. Rigid-body models were calculated by restraining the superfold interfaces described in figure 5D and the covalent disulfide bonds in the EGF stalk and using the SAXS data corresponding to the isoform for which the superfold interfaces are found, i.e. Ten3-A_0_B_0_ for YD-YD, Ten3-A_0_B_1_ for FN-FN and Ten3-A_1_B_1_ for EGF8-ABD. χ^2^ values are calculated by comparing the calculated models against the SAXS data of all isoforms.

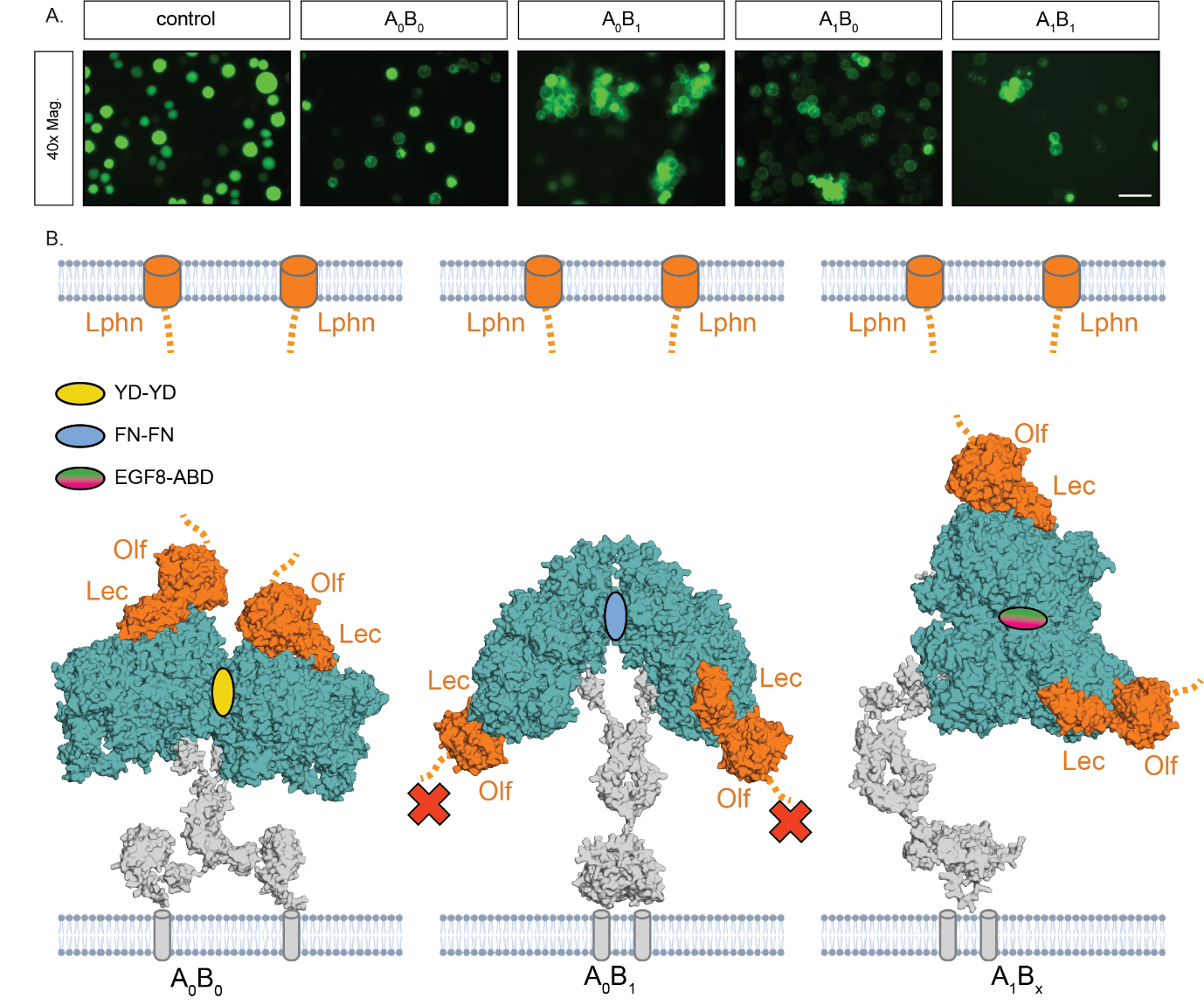

Supplemental Figure 5: Model for isoform-specific teneurin-latrophilin *trans*-cellular complex formation. **A)** Clustering assay of K562 hematopoietic cells electroporated with the mouse Ten3 isoforms at 40x magnification. For all isoforms, GFP signal localises to the membrane in contrast to the soluble GFP that occupies the entire cell area. Scale bar is 100 µm. **B)** Exposure of known latrophilin binding site (Olf-Lec, in orange)^11,23^ for each splicing-dependent compact dimer. In all panels, coloured ellipses indicate the type of intra- and inter-dimeric contacts formed, according to the interface colour codes in Fig. 5D. The rigid-body models of each of the EGF-Ig stalks were calculated using the SAXS data of each isoform (Fig. 5 and Suppl. Fig. 4G).

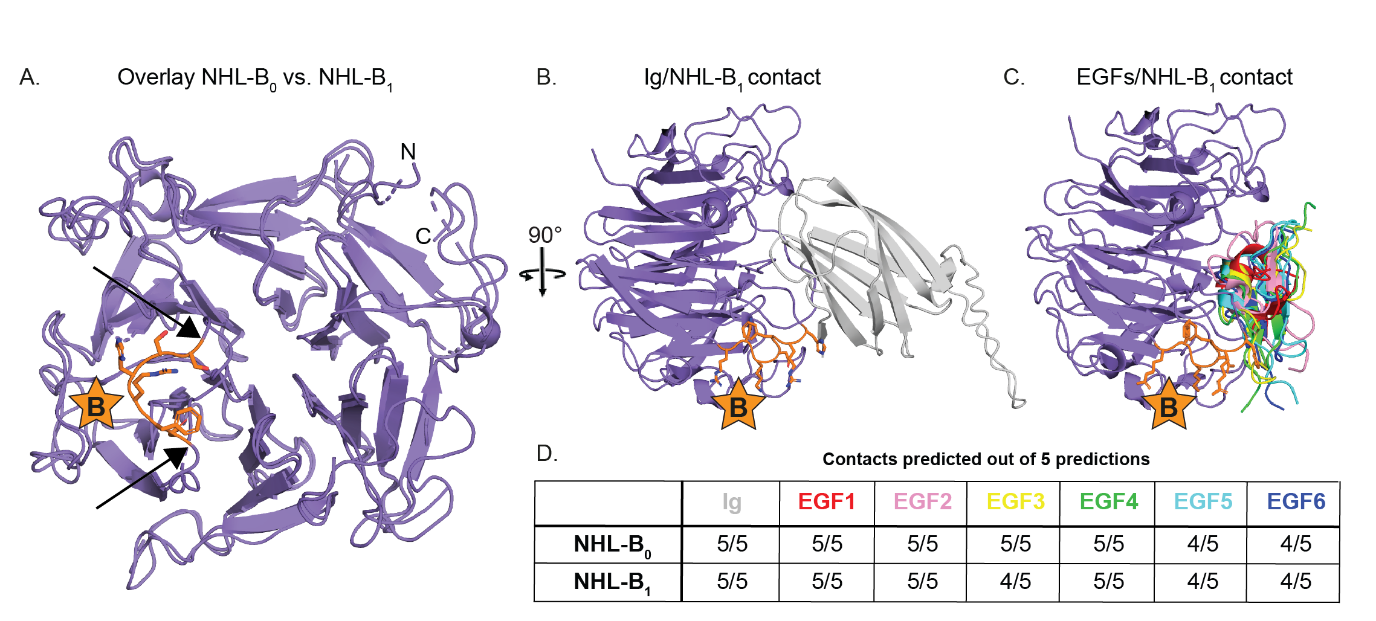

Supplemental Figure 6: Effect of splice insert B on local NHL domain structure and on interactions with EGF and Ig domains. **A)** Structural comparison of NHL domains lacking (B_0_) and containing (B_1_) splice insert B. B_0_ and B_1_ models were built into Ten3-A_0_B_0_ non-compact subunit and A_1_B_1_ compact dimer subunit densities, respectively. Splice insert B is shown as orange stick representation. Boundaries of splice insert B are indicated with arrows. N- (N) and C-termini (C) of the NHL domain are indicated. **B)** Colabfold predictions of the NHL-B_1_ contact with Ig domain (grey). Splice insert B is shown as orange stick representation, and indicated with a star. **C)** Colabfold predictions of the NHL-B_1_ contact with EGF1 through EGF6. Different EGFs predictions have different rainbow colouring, and splice insert B is shown as orange stick representation, and indicated with a star. **D)** Table showing the proportion of predictions containing a direct contact between the EGF or Ig domains and the NHL loop harbouring splice site B - out of 5 total predictions per NHL-EGF or NHL-Ig pair.
